# Dynamic emergence of relational structure network in human brains

**DOI:** 10.1101/2022.05.07.491053

**Authors:** Xiangjuan Ren, Hang Zhang, Huan Luo

## Abstract

Reasoning the hidden relational structure from sequences of events is a crucial ability humans possess, which help them to predict the future and make inferences. Besides simple statistical properties, humans also excel in learning more complex relational networks. Several brain regions are engaged in the process, yet the time-resolved neural implementation of relational structure learning and its behavioral relevance remains unknown. Here human subjects performed a probabilistic sequential prediction task on image sequences generated from a transition graph-like network, with their brain activities recorded using electroencephalography (EEG). We demonstrate the emergence of two key aspects of relational knowledge – lower-order transition probability and higher-order community structure, which arise around 840 msec after image onset and well predict behavioral performance. Furthermore, computational modeling suggests that the formed higher-order community structure, i.e., compressed clusters in the network, could be well characterized by a successor representation operation. Overall, human brains are constantly computing the temporal statistical relationship among discrete inputs, based on which new abstract knowledge could be inferred.

## Introduction

Events in the world are not isolated but are always interconnected according to some hidden structure, which we human beings are constantly pursuing (Pudhiyidath et al., 2021; Schapiro, Rogers, Cordova, Turk-Browne, & Botvinick, 2013). In fact, humans are gifted with tremendous abilities to discover the latent structure behind sequences of events, which is essential for rule inference, future prediction, and decision making (Balaguer, Spiers, Hassabis, & Summerfield, 2016; Garvert, Dolan, & Behrens, 2017; Garvert, Saanum, Schulz, Schuck, & Doeller, 2021; Pudhiyidath et al., 2021; Pudhiyidath, Roomea, Coughlina, Nguyena, & Preston, 2019). For example, after watching several scenes in a movie containing various character combinations, we could quickly form a structural network in mind that signifies connections among characters, such as families, lovers, friends, enemies, colleagues, etc., which are further attached to make a panoramic relational network. Moreover, we could even deduce new knowledge beyond the observations, e.g., reasoning that two seemingly unrelated characters might belong to some common group given their largely overlapping communities.

A principal factor signifying relationships is the transition probability between event occurrences (Aslin, Saffran, & Newport, 1998; Dehaene, Meyniel, Wacongne, Wang, & Pallier, 2015; Ding, Melloni, Zhang, Tian, & Poeppel, 2016; Maheu, Dehaene, & Meyniel, 2019). Greater transition probabilities imply stronger associations, while smaller transition probabilities imply looser links. Previous research has shown that humans, even several-month-old infants, are adept at tracking the transition probability of a series of events (Maheu et al., 2019; Saffran, Aslin, & Newport, 1996) and extracting discrete chunks (Balaguer et al., 2016; Dehaene et al., 2015). Moreover, in addition to the one-dimensional transition chain, recent studies have investigated the learning of more complex networks, such as those with lattice, hexagonal, and community structures (Kahn, Karuza, Vettel, & Bassett, 2018; Kemp & Tenenbaum, 2008; Lynn, Kahn, Nyema, & Bassett, 2020; Mark, Moran, Parr, Kennerley, & Behrens, 2020; Schapiro et al., 2013). Remarkably, humans could not only track transition probabilities, but also infer new abstract (higher-order) statistical structures from event sequences (Garvert et al., 2021; Mark et al., 2020). For example, humans tend to group events into clusters or hierarchical trees, which bias their subsequent reasonings (Balaguer et al., 2016; Pudhiyidath et al., 2021).

Earlier fMRI studies have localized brain regions for the high-level structure representation (Schapiro et al., 2013; Schapiro, Turk-Browne, Norman, & Botvinick, 2016). However, the time-resolved neural implementation of relational structure learning in the human brain – tracking lower-order transition probability and inferring higher-level structure network – and the behavioral relevance, remain elusive. To this end, subjects in the present study performed a prediction task on a sequence of images, with their time-resolved brain activities recorded using electroencephalography (EEG). Unbeknownst to them, the image sequence was generated from a transition network with a community structure that comprises three connected clusters (Fig. 1A). We employed behavioral measurements, time-resolved representational similarity analysis, and computational modeling to seek the neural correlates of structural learning and inference.

**Figure 1.**
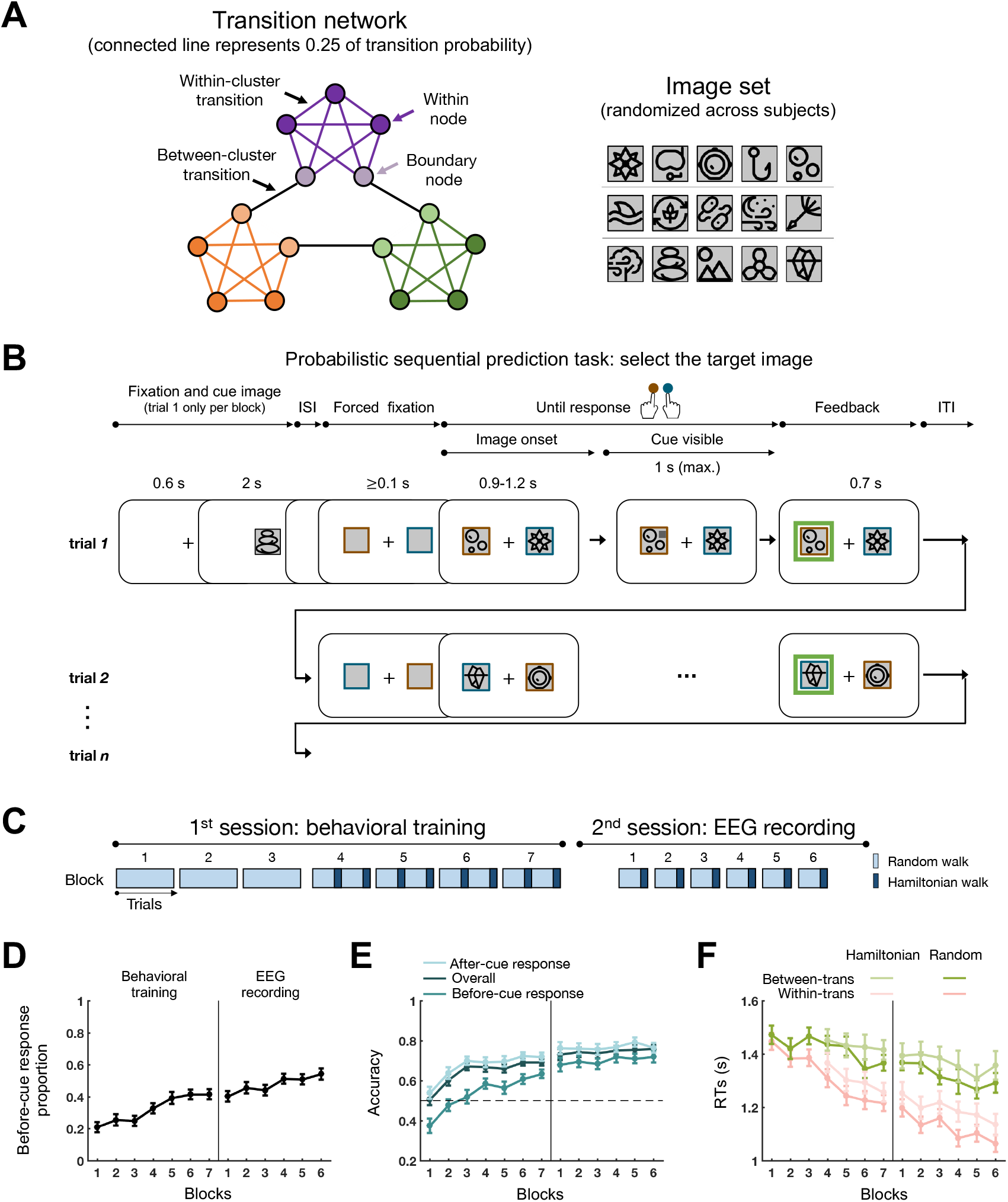
Experimental designs and behavioral results. (A) The underlying transition network with community structure. Each node denotes one image from the image set (right); each connected line denotes one-step transition probability (0.25); boundary nodes (light color) connect clusters; within-nodes (dark color) connect images within the same cluster only. (B) Exemplar block. In the 1st trial, after the initial image, target (directly connected to the initial image, one-step transition) and non-target (non-directly connected) images were bilaterally presented. Subjects chose the target image by pressing corresponding keys according to the border color of the target image using left or right index finger. A cue gradually appeared on target later to hint the correct answer, and subjects were encouraged to make responses before cue. For the rest trials within the block, the target image in each trial served as the initial image of the next trial, and so on. (C) Each subject performed a training session (behavior only) and a formal session with EEG recording on different days. For the last 4 blocks in the 1^st^ session and all blocks in the 2^nd^ session, every 85 random walk trials were followed by 15 Hamiltonian walk trials. (D) Grand average before-cue response proportion for different blocks throughout training and formal sessions. (E) Grand average performance accuracy, for all trials (deep blue), response-before-cue trials (medium blue), and response-after-cue trials (light blue). (F) Grand average RTs for within-cluster (red) and between- cluster transition trials (green), under random-walk (dark) and Hamiltonian-walk (light) paths. Error bars represent ± 1 SEM across subjects.

First, behavioral performances reveal subjects’ successful learning of lower-order statistical relationships and higher-level structure inference. Moreover, the alpha-band neuronal lateralization suggests that subjects shift spatial attention to the target image before making a response, further confirming their structural learning. Most importantly, we demonstrate the time-resolved neural signatures of lower-order transition probability tracking and higher-level structure inference. Specifically, the neural correlates of the transition probability matrix underlying the image sequence emerge around 840 msec after image onset and are further correlated with behavioral performance. Crucially, further analysis of the neural representation of the relational network exhibits its higher-order community structure, i.e., compressed within-cluster distance and enlarged between-cluster distance. Computational modeling advocates that the human brain employs a successor representation strategy to make inferences and form the higher-order community structure from image sequences.

## Results

### Probabilistic sequential prediction task

Thirty-three human subjects performed a sequential prediction task on 15 images (Fig. 1A, right), with the transition probabilities between images determined by a hidden transition network (Fig. 1A, left), unbeknownst to subjects. Each node in the network represents one image, and the lines connecting nodes denote their one-step transition probability. Note that all subjects experienced the same 15 images and transition network, but the correspondence between nodes and images was shuffled across subjects.

As shown in Figure 1B, the 1st trial of each block started with an initial image, followed by two bilaterally presented images (“image onset”) – target and non-target. The target image is directly connected to the initial image in the transition network (one-step transition), while this is not the case for the non-target. Subjects needed to choose the target image as accurately and fast as possible, and both visual and auditory feedbacks were played after responses. Importantly, in the rest of the trials within the block, the target image in each trial served as the initial image in the next trial for subjects to make predictions, and so on. In other words, subjects performed a probabilistic sequential prediction task that spans a succession of nodes that comprise a continuous path within the hidden transition network (Figure 1A, left).

Moreover, to aid the learning process and identify the target, after the “image onset” phase, a cue gradually appeared on the target image to indicate the correct answer (Fig. 1B, “cue visible”) in each trial. Subjects were encouraged to make decisions before the “cue visible” phase if they were confident about their choices. The final reward would depend on the subject’s task accuracy, RT, and eye fixation performance (see more paradigm details in Methods).

### Behavioral performance

Subjects performed two sessions on different days (Fig. 1C): behavioral training (1st session) and formal test with simultaneous EEG recordings (2nd session). The training session also served as a prescreening test to exclude participants who failed to learn the task or keep eye central fixation. All the analyses, except the behavioral part described below, were performed on the 2nd session with EEG recordings.

Behavioral performance improved from training to formal sessions (Fig. 1DE). First, the before-cue response proportion significantly increased (Fig. 1D; LMM1: *F*_(12, 32.72)_ = 10.94, *p* < 0.001) with blocks, indicating that subjects were increasingly more confident to make responses before cue. Second, the behavioral accuracy also increased (Fig. 1E) with blocks, for both before-cue (medium green; LMM2: *F*_(12, 31.67)_ = 8.09, *p* < 0.001) and after-cue (light green; LMM3: *F*_(12, 32.13)_ = 8.37, *p* < 0.001) responses. We further examined whether behavioral performances would be biased by the community structure, as revealed in previous studies (Lynn et al., 2020). Overall, the within-cluster transition showed faster reaction time (RTs) compared to the between-cluster transitions, even after regressing out the practice effect and stimulus recency effect (LMM4: *F*_(1, 32.56)_ = 56.36, *p* < 0.001). Moreover, the advantage of within-cluster over between-cluster held for both Random- (Fig. 1F, dark green vs. dark red; LMM5: *F*_(1, 33.06)_ = 65.33, *p* < 0.001) and Hamiltonian-walk trials (light green vs. light red; LMM6: *F*_(1, 33.03)_ = 42.00, *p* < 0.001), further excluding possible local repetition effects.

Overall, subjects learned both the lower-order transition probabilities and higher-order community structures embedded in the serially presented images.

### Alpha-lateralization spatial attention reveals behaviorally-related transition prediction

Since target and non-target images were presented bilaterally in left and right visual fields, we used the classical alpha-band lateralization neural signature (Ede, Chekroud, Stokes, & Nobre, 2019; Wolff, Jochim, Akyürek, & Stokes, 2017) to examine spatial attention. Specifically, spatial attention would presumably be shifted to the visual field containing the target image, resulting in the corresponding alpha-band lateralization.

Figure 2A plots the grand averaged spatiotemporal lateralization profile (normalized contra- minus ipsi- in reference to target location), for correct trials only (see Methods for details). Around 500 ms after image onset, neural activities in electrodes contralateral to target showed an alpha-band power decrease compared to ipsilateral ones (cluster-based permutation test, left-sided, *p* < 0.01), supporting the attentional allocation to the target side, as hypothesized. Furthermore, the within-cluster transition trials exhibited alpha-band lateralization before cue onset (Fig. 2B, left; cluster-based permutation test, left-sided, *p* < 0.01), while the between-cluster transition trials showed significant attentional lateralization only after cue onset (Fig. 2B, right). The results imply the influence of community structure on spatial attention, consistent with behavioral results (Fig. 1F). Moreover, for within-cluster transitions, the response-before-cue trials (behavioral index) indeed exhibited alpha-band lateralization before cue (neural index; Fig. 2C, left). Interestingly, the response-after-cue trials (behavioral index) still displayed significant yet relatively weaker alpha-band lateralization before cue (Fig. 2C, right). Finally, the before-cue alpha-band lateralization neural index (Fig. 2A) showed positive correlations with RTs (Fig. 2D, left) across all correct trials (Fig. 2D, right), in each subject (one-sample *t*-test, right-sided: *t*(32)=2.48, *p* = 0.009). That is, the stronger attention is allocated to the target side, the faster the prediction speed.

**Figure 2.**
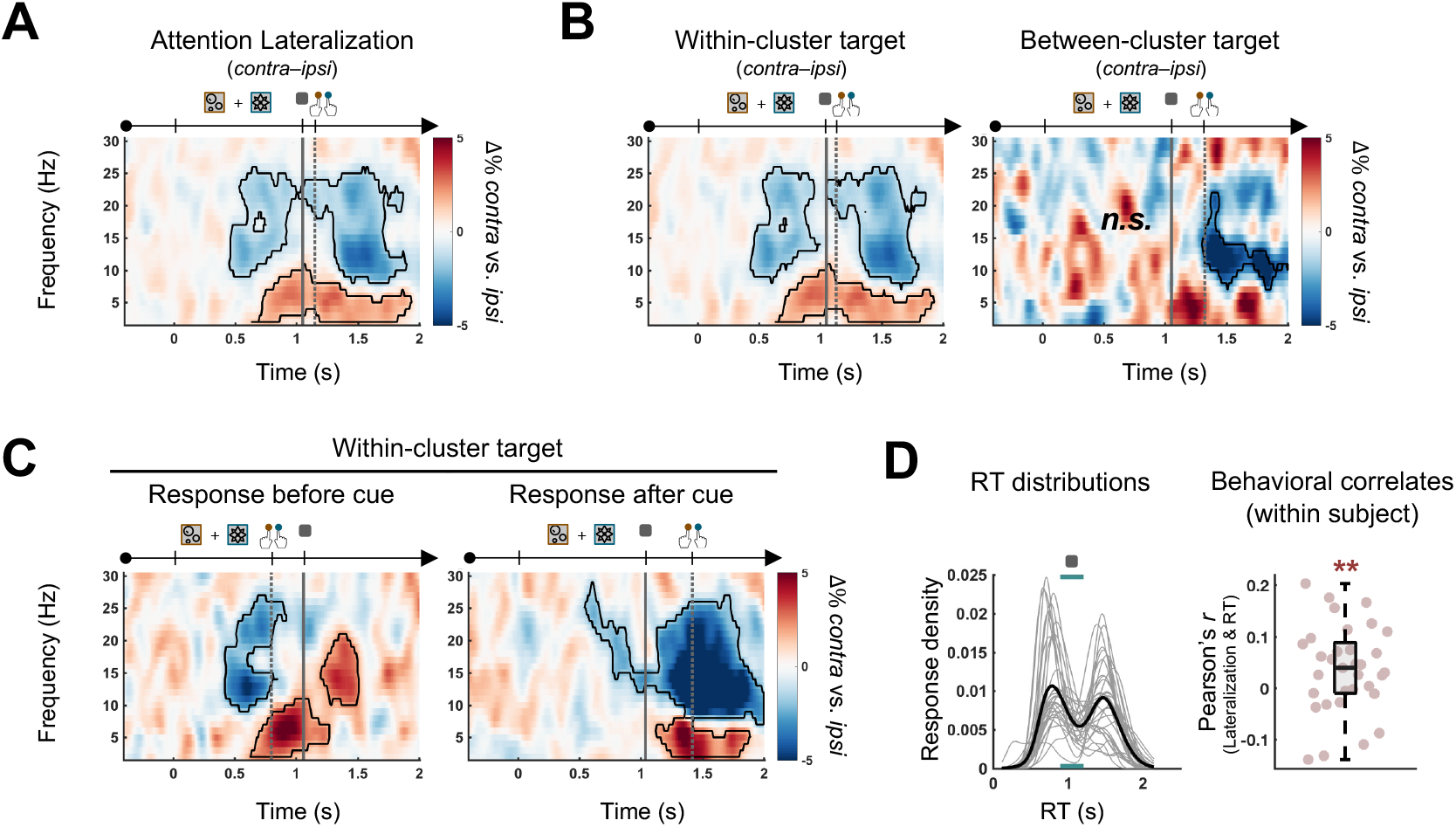
Alpha-band lateralization indexes behaviorally-related spatial attention during transition prediction. (A) Grand average time-frequency representation of contra-ipsi (in reference to target) responses across all correct trials of EEG recordings. Time 0 indicates the onset of target and non-target images. Solid vertical line indicates the cue onset. Dotted vertical line indicates the mean RT. (B) Same as (A) but for within-cluster transition (left) and between-cluster transition trials (right). (C) The same as (A) but for within-cluster transition trials when subjects made responses before cue (left) or after cue (right). (D) Left: RT distribution (all correct trials). Each grey line represents the RT distribution of one subject, which was fitted by a kernel-smoothing distribution. The black line denotes the grand average. The two peaks arise from before- and after-cue responses. The upper grey square represents the cue. The underlying horizontal cyan line denotes the jittered cue onset time. Right: Pearson’s correlation coefficients between averaged before-cue alpha-band neural lateralization index and RTs across all correct trials for each subject. Each dot represents one subject. The middle line in the box plot denotes the median Pearson’s *r* across subjects; Bottom and top lines denote the lower and upper quartiles, respectively; Error bars denote the 99% confidence interval. **: *p*=0.009.

Overall, subjects learned the transition network and shifted spatial attention to the target. Moreover, the attentional neural signature predicts behavior and is also modulated by community structure.

### Neural representation of lower-order transition network and its behavioral relevance

We next examined the neural correlates of the lower-order transition network knowledge subjects had acquired through learning. Each of the 15 images is related to each other via transition probability, which is determined by the corresponding connected nodes within the transition network (Fig. 3A, left). Therefore, the associations between any pair of the 15 images could be characterized by their mutual one-step transition probability (TP), yielding a 15*15 TP design matrix (Fig. 3A, left-top). We further constructed a minimal distance (MD) design matrix with each cell denoting the minimal transition distance for each pair within the transition network (Fig. 3A, left-bottom). Both matrices characterize the transition relationships of the network, among which the TP matrix only represents one-step transition (0 or 0.25 in value), whereas the MD matrix could depict transitions across multiple steps and accordingly entails a more continuous value span.

**Figure 3.**
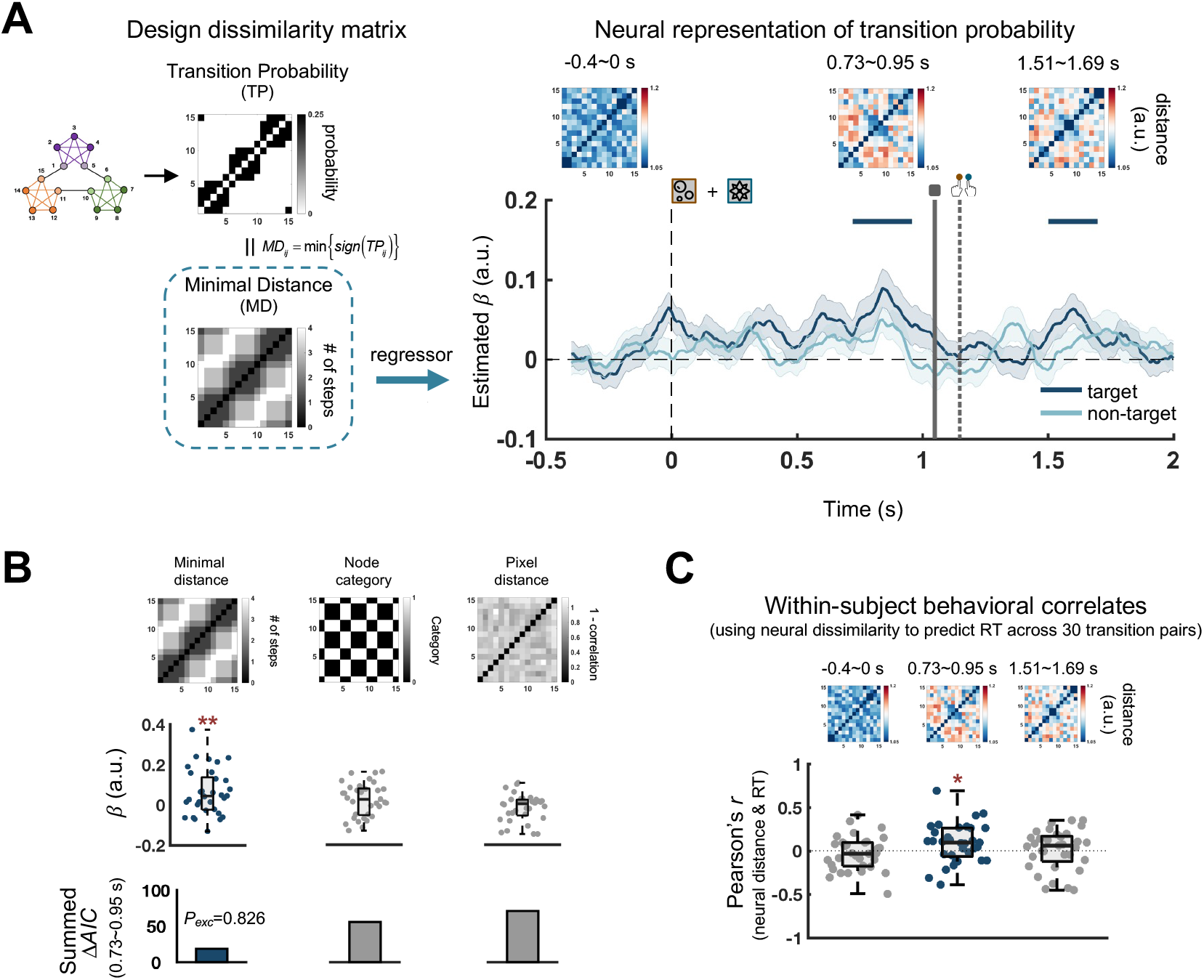
Neural representation of lower-order probability transition between images and its behavioral relevance. (A) Left: Transition probability (TP) design matrix (Upper), with each cell denoting TP (0 or 0.25) for an image pair in the transition network. Minimal distance (MD) design matrix (Lower), with each cell denoting the minimal spanning distance for an image pair in the transition network. Right: Grand average time-resolved GLM results for neural representational dissimilarity defined on target (deep blue) or non-target (light blue) based on MD design matrix. Shadings represent SEM across subjects. Time 0 denotes the onset of target and non-target. Solid vertical line denotes the cue onset. Dotted vertical line denotes the mean RT. Solid horizontal lines denote significant temporal clusters (*p* < 0.01; corrected using cluster-based permutation test, right-sided). Upper small insets plot the neural RDM at different temporal epochs, i.e., before image onset (left; -0.4-0 s), before-cue significant cluster (middle; 0.73-0.95 s), and after-cue significant cluster (right; 1.51-1.69 s). (B) Performance comparisons between MD model, Node category model, and Pixel distance model (upper), based on regression coefficients (middle) and model fitting performances (bottom). For each subject, the model with the lowest AIC was used as a reference to compute ΔAIC for each model. A lower value of Δ AIC summed across subjects indicates better fit. The protected exceedance probability (*P*_exc_) is a metric to evaluate the probability that each specific model outperforms the other models based on the group- level Bayesian model selection. **: *p* < 0.01 (one sample *t*-test, right-sided). (C) Correlation between neural RDM and RTs across 30 image pairs with direct connections in transition network for three temporal windows. Each dot represents one subject. The middle line in the box plot denotes the median Pearson’s *r* across subjects; Bottom and top lines denote the lower and upper quartiles, respectively; Error bars denote the 99% confidence interval. *: *p* < 0.05 (one sample *t*-test, right-sided).

We hypothesize that the neural pattern similarity for image pairs, characterized by neural representational dissimilarity matrix (RDM), would rely on their associations defined in the transition network. Specifically, the closer the two images in the MD matrix (i.e., smaller transition distance), the more similar their neural representations would be (Kriegeskorte, Mur, & Bandettini, 2008). To this end, we ran a general linear model (GLM) on the neural RDM with the MD design matrix as a regressor, at each time and in each subject.

As illustrated in Figure 3A (right panel), there was a significant temporal cluster at 0.73-0.95 sec after image onset (dark blue, cluster-based permutation test, right-sided, *p* < 0.01), indicating the emergence of neural representations of probability transition before cue onset (similar results for the before-cue-response trials; Fig. S1). The significant cluster at 1.51-1.69 sec (cluster-based permutation test, *p* < 0.01), given its appearance after cue, mainly reflects the feedback-based transition learning. As a control, regressing the neural RDM for non-target images using the same MD design matrix did not yield any significant clusters (Fig. 3A, light blue). Furthermore, model comparison showed that design matrixes based on node category or pixel-based distance did not outperform the MD design matrix in characterizing the neural RDM (Fig. 3B) (see results during other time ranges, Fig. S2). Moreover, since the left/right motor response for the target image was balanced across trials, the MD design matrix could not arise from response associations between images, either.

We further tested the behavioral relevance of the neural RDM. We posit that the image pairs with more similar neural representations (i.e., smaller values in neural RDM) would be accompanied by faster behavioral responses since they are associated strongly in the neural space. To this end, we selected all the 30 one-step transition image pairs in the transition network, extracted their respective neural RDM value (Fig. 3C; small insets) at different temporal epochs, and calculated the corresponding RTs when the pair occurred serially during the experiment. A Pearson correlation was then performed between the neural representational dissimilarity and RTs across the 30 pairs, in each subject. As shown in Figure 3C, RTs showed significant correlation to the RDM-based neural index within 0.73-0.95 sec (one-sample *t*-test, right-sided, *t*(32) = 2.21, *p* = 0.017), the time window localized previously (as in Fig. 3A, right), but not for other temporal windows (-0.4-0 s: *t*(32) = –0.89, *p* = 0.810; 1.51-1.69 s: *t*(32) = 0.29, *p* = 0.387).

Overall, the neural representation of the lower-order transition relationship between images emerges around 840 msec after image onset and is further related to prediction behavioral performance.

### Emergence of higher-order community structure: neural evidence and computational modeling

After showing the behaviorally-relevant neural correlates of the lower-order transition network, we next examined whether the neural codes entail the higher-order community structure. Our motivation derives from the advantage of within-cluster over between-cluster transitions shown in both behavior (Fig. 1F) and spatial attention (Fig. 2B), as well as previous findings (Lynn et al., 2020). Moreover, network theories also postulate the emergence of inhomogeneous transition for community structure (Fortunato & Hric, 2016). In other words, instead of exactly tracking the designed lower-order transition probability, the human brain might alter the neural representations in a way that compresses within-cluster and stretches between-cluster distances.

We first compared the neural dissimilarity (i.e., neural RDM from 500 to 900 msec, Fig. 3A and Fig. S1) for image pairs belonging to the same or different clusters. Specifically, as shown in Figure 4A (left), two types of paths that both connect two boundary nodes and span two transitional steps were selected, with one making transitions within the cluster (within-cluster, blue line) while the other across clusters (between-cluster, orange line). We then computed the neural dissimilarities for images at the path endpoints. If the neural distance follows the lower-order transitional matrix, we would expect a similar neural distance for the two paths. Meanwhile, as shown in Figure 4A (right), the within-cluster neural distance was shorter than that of the between- cluster (paired sample *t*-test, two-sided, *t*(32) = –2.783, *p* = 0.009), suggesting the emergence of the higher-order community structure in neural activities (see results during baseline and after cue in Fig. S3).

**Figure 4.**
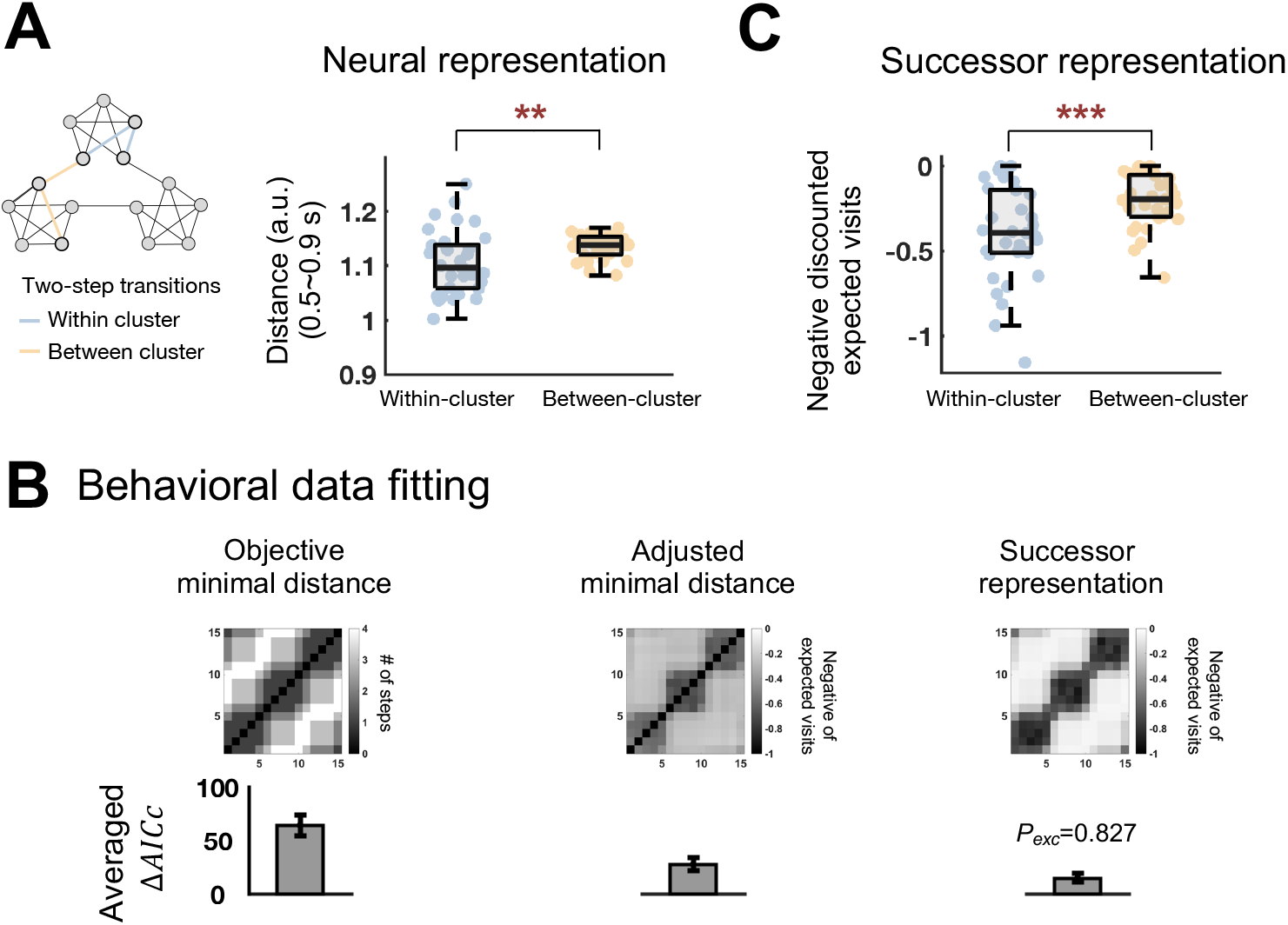
Emergence of higher-order community structure and computational modeling. (A) Left: two types of paths that both connect two boundary nodes and span two steps were selected for comparison to examine the emergence of higher-order community structure. Within-cluster path: transition within cluster (blue line). Between-cluster path: transition across clusters (orange line). Note that they have the same path distance in the lower- level transition matrix. Right: Grand average neural representational distance (i.e., extracted from 500-900 ms neural RDM) for images at the path endpoints for within-cluster (blue) and between-cluster (orange) paths. (B) Three computational models – Objective minimal distance (OMD) (left), Adjusted minimal distance (AMD) (middle), and Successor representation (SR) (right) – were built to fit trial-by-trial behavioral performance per subject. Upper: resulted transitional matrix for the three models. Lower: model comparison in terms of averaged Δ AICc and *P*_exc_ (protected exceedance probability). (C) Grand average distance for within-cluster (blue) and between-cluster (orange) paths in the SR-based transition matrix. Note that both neural (A, right) and behavioral modeling (C) results support the emergence of higher-order community structure. ***: *p* < 0.001, **: 0.001≤ *p* < 0.01 (paired sample *t*-test, two- sided).

To examine the underlying computational mechanism, we built three models – objective minimal-distance model (OMD), adjusted minimal-distance model (AMD), and successor representation model (SR) (Dayan, 1993; Gershman, 2018) – to account for subjects’ trial-by-trial behavioral performances. Specifically, the OMD model simply follows the objective transition network (Fig. 4B, left). The AMD model approximates the transitional relationship between states through iteratively summarizing the recently experienced stimuli (Fig. 4B, middle). The SR model (Fig. 4B, right) assumes a long-term transitional relationship between each state and all its successor states (Dayan, 1993; Gershman, 2018; Stachenfeld, Botvinick, & Gershman, 2017). The three models were then used to fit each subject’s trial-by-trial behavior and their fitting performances were compared based on the summed *ΔAICc* and protected exceedance probability (*P*_exc_) (see details in Methods). As shown in Figure 4B (lower panel), the SR model outperforms the others in characterizing behavioral performance.

Furthermore, the model fitting of behavioral performance also resulted in AMD and SR model’s respective transition matrix (Fig. 4B, upper), from which the within-cluster and between-cluster distance could also be computed to evaluate the higher-order community structure. As shown in Figure 4C, the SR-based transition matrix (the winning model) also displayed the higher-order community structure, i.e., shorter within-cluster distance than between-cluster (paired sample *t*-test, two-sided, *t*(32) = –8.503, *p* < 0.001), thus consistent with the neural findings (Fig. 4A, right).

Taken together, instead of precisely following the objective transition probability, the neural representations launch a new higher-order community structure and form compressed clusters, arising from a successor representation strategy.

## Discussion

Reasoning the hidden relational structure from outside events is a crucial ability we human beings possess to help predict the future and make inferences (Balaguer et al., 2016; Garvert et al., 2021; Pudhiyidath et al., 2021). Here we demonstrate the behaviorally-related neural representations of two key aspects of relational knowledge – lower-order transition probability and higher-order community structure, which occur around 840 msec after image onset, before making a response. Computational modeling suggests that the higher-order community structure arises from a successor representation operation. Taken together, human brains are constantly computing the temporal statistical relationship among discrete inputs, based on which new abstract knowledge is construed.

In spite of limited processing and storage capacity, human beings are good at discovering how components are related to each other, i.e., forming a ‘cognitive map’ about the external world in the mind (Behrens et al., 2018; Bellmund, Gärdenfors, Moser, & Doeller, 2018). Although previously viewed for spatial navigation only, ‘cognitive map’ also mediates non-spatial tasks (Constantinescu, O’Reilly, & Behrens, 2016; Schuck, Cai, Wilson, & Niv, 2016), advocating its general function in relational representations (Nelli, Braun, Dumbalska, Saxe, & Summerfield, 2021; Pudhiyidath et al., 2021). What are the key factors determining the associations characterized in the ‘cognitive map’? The sequence in which events occur, i.e., transition probability, might be a key clue (Lynn et al., 2020; Stachenfeld et al., 2017). Accordingly, statistical learning studies show that humans, even several-month-old infants, can extract statistical transition probabilities and in turn segmenting continuous stimuli (Maheu et al., 2019). This ability might essentially underlie grammar learning in language acquisition (Dehaene et al., 2015). Hippocampus is found to be involved in the process (Schapiro et al., 2013; Schapiro et al., 2016), and interestingly, the hexagonal coding properties of grid cells correspond to the eigenvectors of transition probability matrix in spatial maps (Stachenfeld et al., 2017). Importantly, in addition to the simple one-dimensional transition structure (Ciranka et al., 2022; Liu, Dolan, Kurth-Nelson, & Behrens, 2019; Nelli et al., 2021), events could also be linked in a graph-like network with different structures, e.g., lattice, hexagonal, community, hierarchical tree, etc (Kahn et al., 2018; Kemp & Tenenbaum, 2008; Lynn et al., 2020; Mark et al., 2020; Schapiro et al., 2013). Consistent with previous graph-learning studies (Kahn et al., 2018; Lynn et al., 2020), we demonstrate that human subjects could learn the relationship between images embedded in a transition network. Moreover, the alpha-band lateralization neural signature indicates that subjects shift spatial attention to the target image before making responses, further confirming their acquisition of transition relationships.

Importantly, our results provide new time-resolved evidence for the neural representations of transition probability of a graph-like network. Specifically, images with larger one-step transition probability (i.e., smaller distance in the transition network) come to be represented more similarly in the brain. This neural signature occurs around 840 msec after image onset, before subject make responses, and could well explain the subsequent behavioral performance (i.e., RTs). Previous fMRI studies have revealed multiple brain regions, such as the hippocampus, inferior prefrontal gyrus (IFG), insula, anterior temporal lobe (ATL) in representing the network-based temporal relationship (Pudhiyidath et al., 2021; Schapiro et al., 2013; Schapiro et al., 2016). Thus, the observed neural signature might arise from collective contributions from many brain regions. Although previous studies have also examined how the brain tracks transition probability over time, they mainly focused on expectation violation or event segmentation (Maheu et al., 2019). Here we studied the neural correlates of the transition network denoting a full-scale view of transition probabilities for all image pairs. Moreover, the relatively late emergence of the transitional network in brain activities suggests that relational network learning engages high-level cognitive processing rather than automatic bottom-up schemas.

Most crucially, in addition to unveiling the behaviorally-related neural marker for lower-order temporal statistics, we demonstrate the emergence of higher-order community structures in neural activities. Specifically, neural distances between boundary nodes within the same community become shortened while those between different communities are enlarged, suggesting the formation of compressed clusters in neural representations. Our results are thus commensurate with prior studies revealing that temporal community statistics drive the formation of discrete events (Balaguer et al., 2016), modulate prediction performance (Pudhiyidath et al., 2021), and bias subsequent reasoning, even for non-temporal properties (Pudhiyidath et al., 2019) and in new contexts (Mark et al., 2020). fMRI studies have shown brain regions, including IFG, insula, ATL, STG, etc., which mediate the community structure representation (Schapiro et al., 2013). Moreover, compressed serial replay in spontaneous activities localized in hippocampus is also suggested to mediate new structure inference (Liu et al., 2019).

What is the computational mechanism underlying the formation of a higher-order community structure? Computational models of associative learning and reinforcement learning have been previously used to explain transition probability learning (Ciranka et al., 2022; Maheu et al., 2019), and the successor representation (SR) model could account for the emergence of community structure (Pudhiyidath et al., 2021; Stachenfeld et al., 2017). Here we built three computational models to seek the computational nature of the structural network. We found that the SR model could well characterize the trial-by-trial behavioral performance and most critically, also manifests the higher-order community structure as shown in neural representations, consistent with previous findings (Lynn et al., 2020; Schapiro et al., 2013). Interestingly, although a bit worse, the AMD model also showed reasonably well behavioral fitting performance as well as the higher-order community structure (Fig. S4). The AMD model is akin to the heuristic strategy in decision making, which offers an economic advantage with minimal cognitive cost by relying on simple judgment rules stored in memory (Korn & Bach, 2018).

Relationships between elements usually follow limited structural patterns (Kemp & Tenenbaum, 2008). Therefore, to facilitate learning and inference in new environments, the brain might exploit prototypical structures as schemas on new inputs (Mark et al., 2020). As a result, structure knowledge needs to be coded in a disentangled manner from sensory stimuli, whereby structure and content could be combined in numerous ways. Furthermore, structures could efficiently organize fragmented items in memory, a process that would be beneficial to memory formation (Liu et al., 2019). Recent studies have revealed neural evidence backing the factorized coding of contents and structure in rule learning, working memory, and decision making (Fan, Han, Guo, & Luo, 2021; Kikumoto & Mayr, 2020; Liu et al., 2019). Consistent with these findings, our results also support the dissociated neural underpinnings of structure information – lower-order transition probability and higher-order community structure, which are independent of associated images.

## Materials and Methods

### Experimental Procedures

#### Subjects

Thirty-three human subjects (aged 18-25, 17 female) participated two sessions of the experiment. All of them had normal or corrected-to-normal vision. Subjects completed the behavioral training session and EEG recording session respectively for approximately 110 min and 120 min, receiving 110∼160 and 140∼210 RMB for their time. The study had been approved by the Institutional Review Board of School of Psychological and Cognitive Sciences at Peking University (#2019-02-08). Subjects provided written informed consent in accordance with the Declaration of Helsinki.

#### Apparatus and stimuli

Subjects were seated approximately 100 cm (*i.e.*, 1.5 cm≍ 1 ^°^) in front of a 28-inch display++ monitor (60.6 × 38.1 cm, 1,680 1,050 pixels, 100-Hz refresh rate). The display of stimuli and recording of responses were controlled by a Dell computer using Matlab R2018b and PsychToolbox-3 (Brainard, 1997; Pelli, 1997). The stimuli were 15 cartoon images with abstract semantics (https://www.flaticon.com/packs). The image sequence was generated according to a transition network (see the following Design). Subjects needed to keep eye gaze over the fixation cross throughout the whole trial, and the eye movements of their dominant eyes were continuously monitored by an eye tracker (Eyelink-1000, SR Research Ltd.).

#### Transition network

Unbeknown to the subjects, we used a community-structured transition network with 15 nodes to generate the target image sequences across trials per block, with each node denoting one image in the image set (Fig. 1A). Each node had an even transition probability of 0.25 to transition into one of four other nodes. The network consists of three interconnected communities of five nodes. In each community, three nodes (“within nodes”) were fully connected to the other four nodes within the same community and the remaining two nodes (“boundary nodes”) were connected to the three within nodes and one node from other communities. All subjects experienced the same 15 images and a same transition network, but the correspondence between nodes and images was shuffled across subjects.

#### Transition prediction task

Please see Figure 1B for the temporal course of one trial. The 1^st^ trial of each block started with an initial image, followed by the presentation of target and non-target images. The target image was directly connected to the initial image in the transition network (one-step transition), while the non-target image was not directly connected to the initial image. In the rest trials within the block, the target image in each trial served as the initial image in the next trial for subjects to make predictions, and so on. The order of target images across trials was prescribed by either a Random or a Hamiltonian walk on the transition network. A Random walk is a path across a transition network created by taking repeated random steps between any two directly connected nodes, while a Hamiltonian walk is a path that visits each node exactly once in the consecutive 15 trials (Newman, 2010).

#### Experimental Procedure

In each experimental block, the 1^st^ trial started with the presentation of an initial image for 2 s followed by a fixation screen. For the rest trials within the same block, each trial started with a fixation screen containing two gray boxes (3 ^o^ x 3 ^°^) presented bilaterally (5 ^°^from fixation). At the beginning of each trial, only when their eye fixation lasted for more than 0.1 s, the target and non-target images would appear bilaterally in the two gray quadrates, and then subjects selected the target image as quickly and accurately as possible. Before response within a trial, subjects’ gaze positions of dominant eyes were continuously monitored. Larger deviation from the fixation cross would trigger a sound warning. Notably, to remove fixed correspondence between motor response and spatial location, the border color of the target and non-target images were randomly set in either orange or cyan per trial, and subjects chose the target by pressing keys (left index finger for ‘z’; right index finger for ‘m’) with the same color as that for border of the target box (i.e., ‘z’ for orange and ‘m’ for cyan, or vice versa).

To facilitate the transition probability learning, after 0.9–1.2 s presentations of target and non-target images, a cue (a 0.4 ^°^ x 0.4 ^°^ gray square) appeared gradually on the target image to indicate to subjects the correct answer. Note that subjects were encouraged to make response before cue onset if they were confident. Once subjects made responses, either before or after cue and either correct or incorrect, the target image would be highlighted by a green frame. Meanwhile, an auditory feedback was also presented indicating whether the response was correct, incorrect or time-out. The feedback screen lasted for 0.7 s. During the EEG recording, the inter-stimulus interval (ISI) and inter-trial interval (ITI) were both selected from a uniform distribution ranging from 1.5 to 2 s.

Subjects received rewards based on their behavioral performances and eye fixation. Specifically, correct response before cue would be 100 virtual coins. Otherwise, the accessible coins decreased linearly with time, declining from 100 to 0 within 0.3 s and from 0 to -50 for the remaining 0.7 s. In other words, before- and after-cue responses yielded higher and lower rewards, respectively. Moreover, incorrect (including time-out) responses or failure eye fixation yielded negative rewards, i.e., -50 and -100 coins, respectively. Therefore, in order to accumulate larger rewards, subjects need to eliminate erroneous selection and fixation violations, and at the same time make their decision as quickly as possible.

#### Training and formal sessions

Each subject completed two sessions—training session (behavior only) and formal session (with EEG recordings)—on different days. In the training session, subjects accomplished 1,500 trials, and the sequence consisted of 700 Random walk trials (i.e., transition at each node was uniformly selected from its four connections), followed by eight repeats of 85 Random walk trials and 15 Hamiltonian walk trials (i.e., every node was visited exactly once). The first 700 and 800 trials were binned into three and four blocks respectively. Subjects were not informed about the trial types. Moreover, to eliminate the expectation difference between boundary-boundary transition across communities and boundary- within transition within a community in the Hamiltonian walk trials, we randomly selected one fixed Hamiltonian path for each subject as previous studies (Schapiro et al., 2013). In the training session, one of the subjects only completed 5 blocks (1,100 trials).

Subjects who successfully controlled eye fixations and performed reasonably well in the training session would be invited to participate in the formal session with EEG recordings. In the formal session, subjects performed the same task with EEG recordings, completing six blocks of Random and Hamiltonian walk trials. Note that the correspondence between image and node in the network was fixed for each subject in two sessions, which means subjects did not need to relearn the transition relationship between images in the formal session.

Following the EEG recording, subjects completed a questionnaire that assessed their learning about the transition network. The question was: “Did you find the rules underlying the image sequences? When did you detect the rules? Please briefly describe what you have learned.” About 60% and 25% subjects mentioned that the target image sequences were divided into three groups and certain images acted as a pivot connecting two groups.

### Behavioral Analysis

#### Response ratio, accuracy, and RT

The trials were divided based on whether subjects made response before (before-cue response trials) or after cue onset (after-cue response trials). We calculated accuracy as their correct response proportions for each type of trials. Correct responses were defined as those with correct button press and successful eye fixation. RTs were defined based on the stimulus onset and were computed for correct trials only.

#### Computational modeling

Three computational models – Objective minimal distance model (OMD model), Adjusted minimal distance model (AMD model), and Successor representation model (SR model) (Dayan, 1993; Gershman, 2018) – were used to characterize the trial-by-trial behavioral performance in each subject.

##### OMD model

The OMD model simply follows the objective transition network, in which the expected visits (or relational strength) between node pairs are defined as reciprocal of their minimal distance.

##### AMD model

The AMD model is a kind of OMD-based heuristics, which approximates the expected visits of transitions from node *s* to node *s*′ through summarizing the recently experienced stimuli within a memory constraint window. Specifically, the AMD assumes subjects’ memory capacity is finite and hence they could only remember a limited numbers of stimuli recently appeared. In each trial, the to be counted stimuli are those falling in a window with fixed-length (*T_mw_*) that stretch backward from the current trial. The expected visits of a transition pair are calculated as the proportion of transitions between *s* and *s*′ (direction-irrelevant) in their overall occurrences:

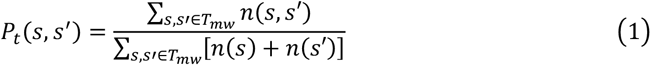

where *n*(*s*) and *n*(*s*′) denote the number of occurrences of stimulus *s* and *s*′ in the current memory window, *T_mw_*; n(*s*, *s*′) is the corresponding number of occurrences of transition from *s* to *s*′ and its reverse, regardless of whether the transition is one-step; *T_mw_* is a free parameter denoting the memory constraint (*T_mw_* > 2), in the unit of trial lengths.

The memory window slides along with trials, during which the proportions for a transition pair are iteratively summed and then normalized in each trial:

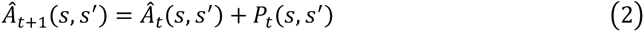

where *Â* is the internal representation of the transition matrix updated through AMD model; the subscripts of *Â* denote trial *t* and *t*+1. For both OMD model and AMD model, the diagonal in the symmetric matrix is nonsense.

##### SR model

The SR model assumes a long-term transition relationship between one state and all its successor states. Mathematically, the SR is a *n* × *n* matrix, with *n* denoting 15 images in the experiment. Each cell of the SR matrix represents the expected discounted visits from one node (or state, *s*) to another node (*s’*), defined as:

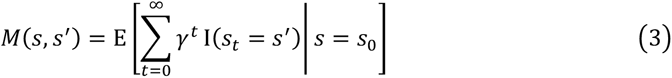

where *γ* is a discount factor (0 < *γ* < 1), *s*_#_ denotes the state transitioned from state *s*_-_ after *t* time-steps, and Ι(·) is a one-hot vector, with its value set to 1 if the successor state is *s*′.

The SR model was learned and updated using the temporal difference learning (Sutton & Barto, 2018), after each trial, by:

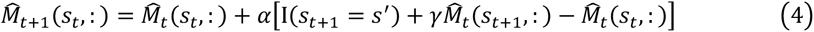

where 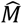 is the internal representation of the transition matrix updated through SR model; *α* is the learning rate (0 < *α* < 1). Specifically, for any transitions from *s*_t_ in trial *t* to *s*_t+1_ in trial *t*+1, the successor matrix (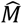) would be updated based on the temporal difference error in the square bracket.

##### Choice

We assumed that subjects made a choice based on the expected visits (or value) difference between the two options (target and non-target). To account for the stochasticity in subjects’ choices, we assumed that subjects’ estimation of the value difference (target minus non-target, *V_T_* – *V_D_*) was contaminated by a Gaussian noise: *ε*∼*N*(0, 1). Whether subjects responded before cue onset depended on whether the value of *V_T_* – *V_D_* + *ε* was greater than a decision threshold (*L*). As the consequence, the probability for each kind of choice (correct response before cue, incorrect response before cue, or response after cue) was defined as:

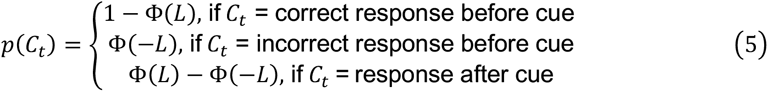

where Φ is the Gaussian CDF with μ = *V_T_* – *V_D_*. and *σ* = 1. Subjects’ choice on each trial was modeled as a multinomial random variable determined by *p*(*C*_t_).

##### Model fitting and comparison

For each subject, we fit the models to subjects’ choice in each trial using maximum likelihood estimates (MLE). Given the choice in each trial has a choice probability, *p*(*C*_t_), the likelihood function derived from multinomial distribution was used to describe the relationship between subjects’ responses and the model predictions. The *fminsearchbnd* (J. D’Errico), a function based on *fminsearch* in MATLAB (MathWorks), was used to search for the parameters that minimized negative log likelihood. To verify that we had found the global minimum, we repeated the searching process for 100 times with different starting points. We compared the goodness of fit of each model based on the Akaike information criterion with a correction for sample sizes (AICc) (Akaike, 1974; Hurvich & Tsai, 1989) and group-level Bayesian model selection (Daunizeau, Adam, & Rigoux, 2014; Rigoux, Stephan, Friston, & Daunizeau, 2014; Stephan, Penny, Daunizeau, Moran, & Friston, 2009). Note that the AICc values punishes number of free parameters, and lower AICc values indicate better fitting performance. Specifically, in each subject, the model with lowest AICc was used as a reference to compute ΔAICc for all models. In addition to the ΔAICc, we also used the group-level Bayesian model selection to compute the protected exceedance probability (*P*_exc_), a metric to evaluate the probability that each specific model outperforms the other models.

#### Transition matrix of model fitting

All three models were used to fit each subject’s trial-by-trial behavior, respectively. In total, OMD, AMD and SR model have one (*L*), two (*T_mw_* and *L*) and three (*γ*, *α*, and *L*) parameters, respectively. The internal transition matrix of AMD and SR model (Fig. 4B, upper panels) were extracted from individual subject based on the best parameters estimated using MLE (see model fitting) and then averaged across subjects.

### EEG analysis

#### Data acquisition and Preprocessing

The EEG signals were acquired using a 64-channel EasyCap. Data was recorded at 500 Hz using two BrainAmp amplifiers (Brain Products) and BrainVision Recorder software (Brain Products), with one additional electrode placed below the right eye as vertical electro-oculograms (EOG). The online EEG recordings were sampled at a rate of 500 Hz. During data acquisition, we used FCz and Cz respectively as reference and ground electrodes, whose impedances were reduced to below 5 *k*Ω . All other electrode impedances were kept below 20 *k*Ω.

Standard preprocessing was performed using Matlab R2016b and the FieldTrip package (Oostenveld, Fries, Maris, & Schoffelen, 2011). Data were band-pass filtered between 0.5 and 45 Hz and then segmented into epochs of 3.1 s relative to stimulus onsets (-0.6 s before and 2.5 s after the image onset). Trials with extremely high noises by visual inspection were manually excluded. Bad channel with abnormal voltage was interpolated with the weighted average of its neighboring electrodes. Independent component analysis was performed to remove eye movement and other artifactual components and the remaining components were then back-projected to the EEG channel space. No subject was excluded for excessive head movement or other artifacts. Only trials with correct button press and satisfactory eye fixation were used for subsequent analyses.

#### Lateralization analysis and behavioral relevance

Time-frequency analysis was performed based on the multi-taper time-frequency transformation of Hanning-window-tapered data. We estimated spectral power at frequencies between 2 and 40 Hz in 1-Hz step, using a fixed 300-ms sliding window that was applied to each trial in 10-ms steps from -0.6 s to 2.5 s relative to image onset. We calculated the lateralization index (Ede et al., 2019) to denote spatial attention, as:

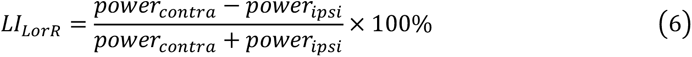

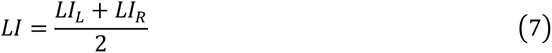

*power_came_* and *power_ipsi_* denote the power in electrode cluster contralateral or ipsilateral to the target side in each trial. Electrodes in spatial lateralization analysis were defined based on previous studies (Ede et al., 2019; Wolff et al., 2017): O1, PO3 and PO7 for left, O2, PO4, PO8 for right.

To examine the behavioral relevance of attentional lateralization index, in each subject, the alpha-band lateralization index within time and frequency clusters that showed statistical significance before cue onset were calculated for each trial, and a Pearson’s correlation analysis was then computed between the lateralization indexes and the corresponding RTs across trials, in each subject. Note that only correct trials were included.

#### Representational Similarity Analysis (RSA) and behavioral relevance

The preprocessed EEG data was first *z*-scored over all trials, for each electrode and at each time point. To obtain the neural representational dissimilarity matrix (RDM), we first averaged trials whose target image was the same and presented at the same location (left or right) and then calculated the Euclidean (1 – correlation) distance between the whole-scalp neural signals of each image pair, resulting a 15 × 15 matrix at each time point and in each subject. The procedure was performed for left and right side presentation separately and the two results were then averaged, yielding the neural RDM at each time point per subject. The neural RDMs were further smoothed over time by convolution with a 60 ms uniform kernel (Luyckx, Nili, Spitzer, & Summerfield, 2019).

A 15 × 15 transition probability (TP) matrix and a minimal distance (MD) matrix were used as the model RDMs to examine the neural RDM. For the TP design matrix, each cell denotes the one-step transition probability for the image pair in the transition network. For the MD matrix, each cell denotes the minimal transition distance (step) for each pair within the transition network. Notably, the TP matrix only represents one-step transition (0 or 0.25 in value), whereas the MD matrix could depict transitions across multiple steps and thus entails a more continuous value span.

We hypothesize that the neural pattern similarity for image pairs, characterized by neural representational dissimilarity matrix (RDM), would rely on their associations defined in the transition network. Specifically, the closer the two images in the MD matrix (i.e., smaller transition distance), the more similar their neural representations would be. To this end, we ran a general linear model (GLM) on the neural RDM with the MD design matrix as a regressor, at each time and in each subject. Note that since neural RDM and model RDM were both symmetrical, only the lower triangles were used for further linear regression analysis.

As a control analysis, we performed the same GLM analysis but used the neural RDM defined by non-target stimuli. Moreover, model RDMs based on two additional models (node category and pixel-based distance) were tested. For the node category model, each cell of the RDM represents whether the row and column nodes belong to the boundary (‘1’) or within (‘0’) node. For the pixel-based model, each cell represents the distance between the row and column nodes (images) in terms of pixel similarity (1 – Pearson’s correlation coefficient).

Given the linear relationship between AIC and mean squared error in linear regressions, we could convert residuals into AIC for each model, at each time point, and for each subject. When evaluating the goodness-of-fit, we used the Δ AIC and protected exceedance probability (*P*_exc_) based on the AIC or log-evidence averaged in the time window of 0.73-0.95 s (see cluster-based permutation test for details).

To test the behavioral relevance of the neural RDM, we selected all the 30 one-step transition image pairs in the transition network, extracted their respective neural RDM value averaged within three temporal epochs (see below), and calculated the corresponding RTs when the pair occurred serially during the experiment. A Pearson correlation was then performed between the neural RDM value and RTs across the 30 pairs, in each subject. Three temporal epochs were selected based on group results: before image onset (-0.4∼0 s), before-cue significant cluster (0.73∼0.95 s), and after-cue significant cluster (1.51∼1.69 s).

### Statistical Analysis

Standard statistical tests, such as paired-sample *t*-test, were used to assess the statistical significance at the group level. Linear mixed models (LMM) were estimated using the ‘fitlme’ function in Matlab R2016b, with *F* statistics, degree of freedom of residuals (denominators), and *P*-values approximated by Satterthwaite method (Fai & Cornelius, 1996; Giesbrecht & Burns, 1985; Kuznetsova, Brockhoff, & Christensen, 2017). Specifications of random effects in LMMs were kept as maximal as possible (Barr, Levy, Scheepers, & Tily, 2013) but without overparameterizing (Bates, Kliegl, Vasishth, & Baayen, 2015).

LMM1, LMM2 and LMM3: the before-cue response proportion, accuracy for before-cue responses and accuracy for after-cue responses are, respectively, the dependent variables; fixed effects include an intercept and the main effect of categorical variable blocks; random effects include random slopes of blocks within subjects and random subject intercept.

LMM4: the RTs of all correct trials are the dependent variables; fixed effects include an intercept, the main effects of trial number (the practice effect; log-transformation), the recency (defined as the number of trials since the last instance of the current target image), the walk types (Random- versus Hamiltonian-walk), the distractor types (same versus different cluster as the target) and the transition types (within- versus between-cluster); random effects include the corresponding random slopes of the main fixed effects within subjects and random subject intercept.

LMM5 and LMM6: the RTs in Random- and Hamiltonian-walk trials are the dependent variables respectively; fixed effects include an intercept, the main effects of categorical variable blocks and transition types (within- versus between-cluster); random effects include random slopes of blocks and transition types within subjects, and random subject intercept.

Moreover, we identified time or time-frequency windows that had above-chance activation using cluster-based permutation tests (Maris & Oostenveld, 2007). Specifically, for the RSA analysis, adjacent time lags with significantly positive decoding performance at the uncorrected significance level of .05 by right-sided one-sample *t* tests were grouped into clusters and the summed *t*-value across the time lags in a cluster was defined as the cluster-level statistic. We randomly shuffled the target labels across trials and performed the time-resolved RSA analysis on the shuffled data from which the maximum cluster-level statistic was computed. This procedure was repeated for 100 times to produce a reference distribution of chance-level maximum cluster-level statistic, based on which we calculated the *p* value for each cluster in real data.

## Author contributions

X.R., H.Z., H.L. designed the experiment, X.R. performed the experiment and data analysis. X.R., H.Z., H.L. wrote the paper.

## Acknowledgments

X.R. was supported in part by the Postdoctoral Fellowship of Peking-Tsinghua Center for Life Sciences. H.Z. was supported by National Natural Science Foundation of China Grants 31871101 and 32171095, and Peking-Tsinghua Center for Life Sciences. H.L. was supported by the National Science and Technology Innovation 2030 Major Program 2021ZD0204103, and National Natural Science Foundation of China Grants 31930052.

## Supplementary materials

**Figure S1.**
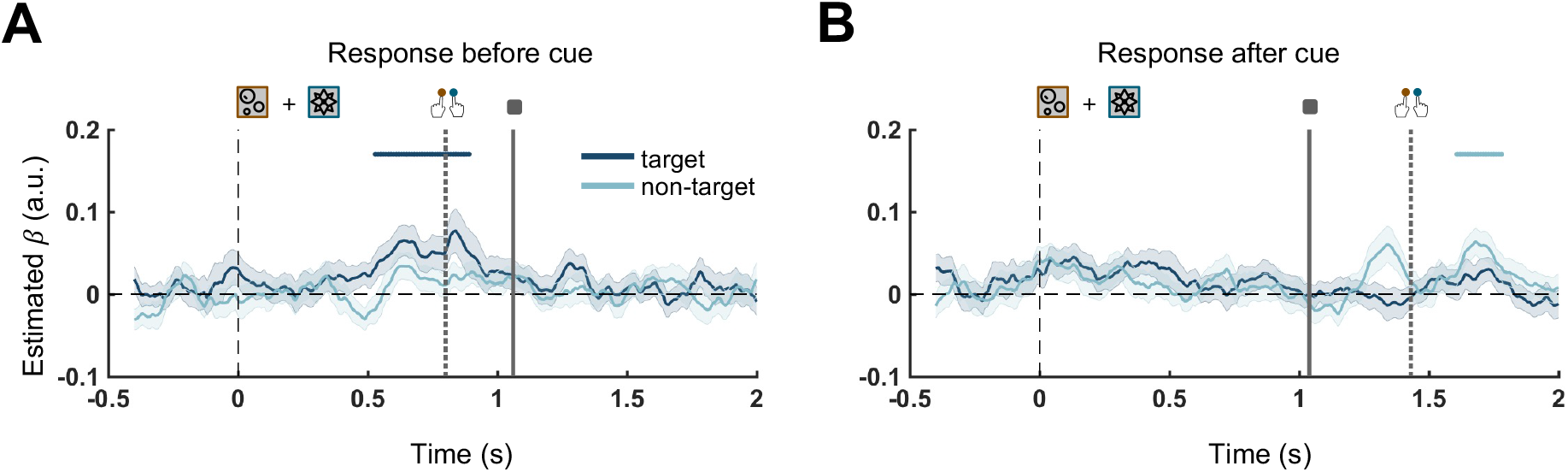
Decoding results for response-before-cue (A) and response-after-cue trials (B). Shadings represent SEM across subjects. Time 0 denotes the onset of target and non-target. Solid and dotted vertical lines denote the cue onset and the mean RT respectively. Solid horizontal lines above the curves denote significant temporal clusters, which are 0.53-0.89 s and 1.61-1.78 s separately for (A) and (B) (*p* < 0.05; corrected using cluster-based permutation test, right-sided).

**Figure S2.**
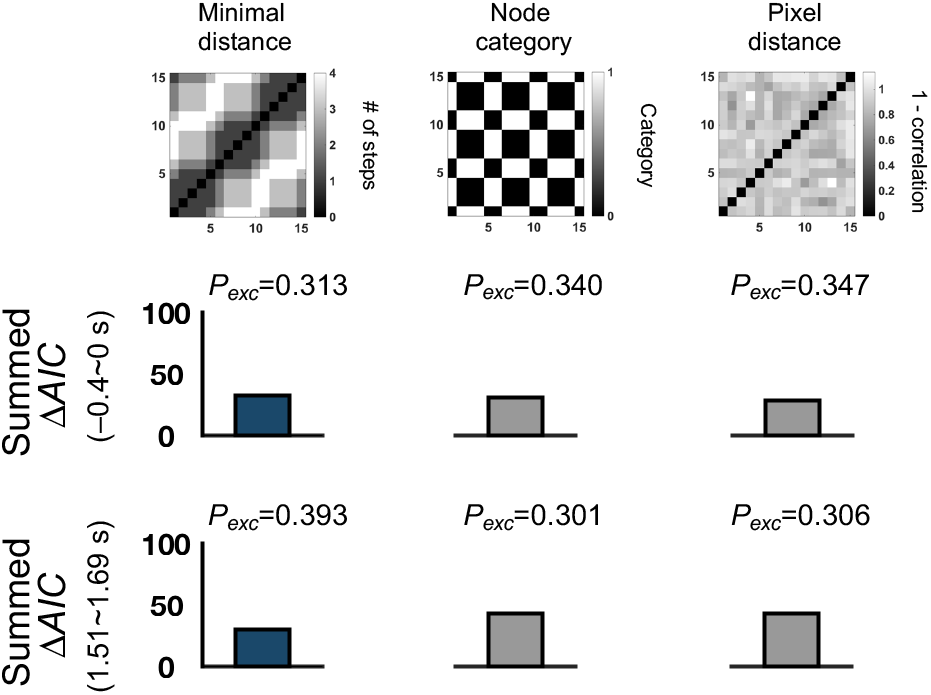
Model comparisons for baseline (middle) and after-cue time range (bottom). For each subject, the model with the lowest AIC was used as a reference to compute ΔAIC for each model. A lower value of ΔAIC summed across subjects indicates better fit. The protected exceedance probability (*P*_exc_) is a metric to evaluate the probability that each specific model outperforms the other models based on the group-level Bayesian model selection.

**Figure S3.**
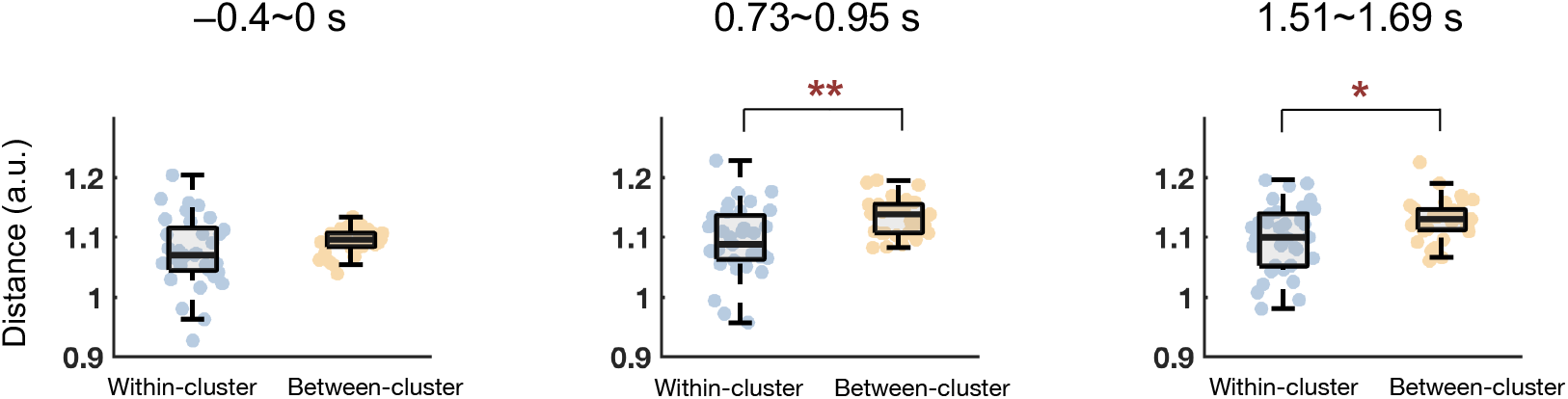
Grand average neural representational distance for images at the path endpoints for within-cluster (blue) and between-cluster (orange) paths. Left: -0.4∼0ss; middle: 0.73∼0.95 s; right: 1.51∼1.69 s. **: *p* < 0.01, *: 0.01 ≤ *p* < 0.05 (paired sample *t*-test, two-sided).

**Figure S4.**
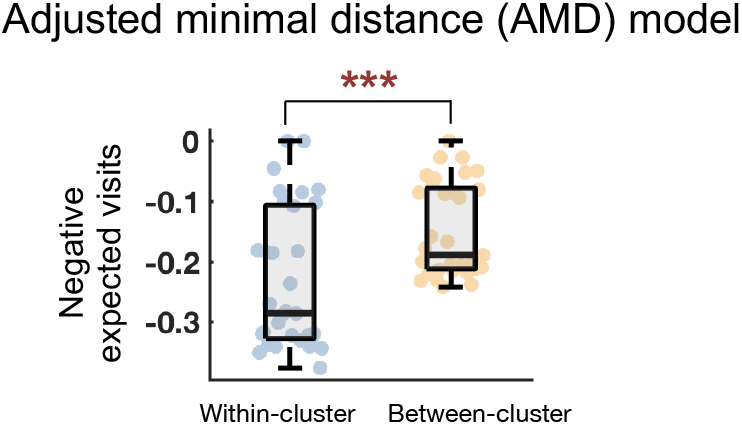
Grand average distance for within-cluster (blue) and between-cluster (orange) paths in the AMD model-based transition matrix. ***: *p* < 0.001 (paired sample *t*-test, two-sided).

